# *CLOCI:* Unveiling cryptic gene clusters with generalized detection

**DOI:** 10.1101/2023.06.20.545441

**Authors:** Zachary Konkel, Laura Kubatko, Jason C. Slot

## Abstract

Gene clusters are genomic loci that contain multiple genes that are functionally and genetically linked. Gene clusters collectively encode diverse functions, including small molecule biosynthesis, nutrient assimilation, metabolite degradation, and production of proteins essential for growth and development. Identifying gene clusters is a powerful tool for small molecule discovery and provides insight into the ecology and evolution of organisms. Current detection algorithms focus on canonical “core” biosynthetic functions many gene clusters encode, while overlooking uncommon or unknown cluster classes. These overlooked clusters are a potential source of novel natural products and comprise an untold portion of overall gene cluster repertoires. Unbiased, *function-agnostic* detection algorithms therefore provide an opportunity to reveal novel classes of gene clusters and more precisely define genome organization. We present *CLOCI* (Co-occurrence Locus and Orthologous Cluster Identifier), an algorithm that identifies gene clusters using multiple proxies of selection for coordinated gene evolution. Our approach generalizes gene cluster detection and gene cluster family circumscription, improves detection of multiple known functional classes, and unveils noncanonical gene clusters. *CLOCI* is suitable for genome-enabled small molecule mining, and presents an easily tunable approach for delineating gene cluster families and homologous loci.

## INTRODUCTION

Gene clusters are chromosomal loci that contain two or more adjacent genes that are co-inherited with cooperative biological functions. Gene clusters are found across the tree of life and encode diverse metabolic and ecological phenotypes (1–6). Perhaps the most well-studied category of gene clusters is biosynthetic gene clusters, which produce small molecule “secondary/specialized metabolites” that mediate biotic and abiotic interactions. Specialized metabolites have diverse activities with applications in industry and agriculture (7) and a prominent source of new drugs in healthcare (8). These compounds include UV-absorbing carotenoids (9), antimicrobial compounds (10,11), and siderophores (12,13). Beyond specialized metabolite biosynthesis, gene clusters can encode pathways that degrade antagonistic compounds (14), catabolize carbon sources and amino acids (15–17), assimilate nutrients (18), or produce essential vitamins (19) and proteins (20–22). Gene clusters can be thought to span a continuum from generalized to specialized based on how broadly applicable the cluster is to organisms’ niches. Generalized metabolic cluster repertoires present a window into essential metabolic processes (23,24) and widely conserved ecological functions (19,25–27), while specialized cluster repertoires provide insight into ecological niche (28–31).

Gene cluster detection is a transformative tool for specialized metabolite drug discovery in the genomic era. Fungal drug discovery rapidly introduced novel drugs in the 20^th^ century by implementing laborious untargeted chemical screening, though the field is increasingly hampered by redundant compound identification due to a focus on easily cultivable biodiversity (32). Additionally, some specialized metabolites are not produced under standard laboratory conditions and myriad expression conditions are often tested to induce silent pathways (33). A reservoir of novel specialized metabolites thus remains hidden in organisms and biosynthetic pathways that are elusive under laboratory conditions (34).

Genomic gene cluster detection unveils specialized metabolite biosynthesis pathways from difficult-to-culture organisms and repressed biosynthetic pathways. Gene cluster detection algorithms have unveiled biosynthetic pathways with novel specialized metabolites (35) and identified novel specialized metabolic pathways that are epigenetically repressed (36). Once gene clusters are identified, it is then possible to predict gene functions and systematically characterize them by heterologous expression (37).

The most widely implemented biosynthetic gene cluster detection algorithms are function-centric (1,38). Function-centric algorithms search for profile models of “core” biosynthetic proteins commonly associated with described biosynthetic gene clusters of bacteria and filamentous Ascomycota (Fungi). Greater than 94% of the fungal clusters in the Minimum Information about Biosynthetic Gene Clusters (MIBiG) database (39) originate from the filamentous fungi in the phylum Ascomycota (Supplementary Figure 2), leading to bias in fungal gene cluster models used for detection. Fungal profile models are largely derived from the “canonical” core genes of polyketide, nonribosomal peptide, and terpene biosynthetic clusters of Ascomycota (40). Function-centric detection is thus expected to identify clusters in the lineages the models are derived from, though it may have poor precision in lineages with “noncanonical” specialized metabolite gene clusters that lack core genes upon which the models rely (34,41–45). To account for noncanonical classes, the function-centric software *antiSMASH* incrementally expands its scope by accumulating models of novel biosynthetic gene cluster core functions (1). Each update increases the breadth of future searches to include new biosynthetic classes, though the function-centric paradigm cannot infer novel noncanonical gene cluster classes *de novo*.

Some gene cluster detection algorithms are function-agnostic, relying instead on using general properties of gene clusters to detect them. Two function-agnostic methods are direct analysis of gene co-expression (45,46) and modeling coordinated expression through identification of shared intergenic motifs (47). Co-expression detection predicts gene clusters from transcriptomic data by identifying significantly coexpressed genes within the same locus. In practice, unsupervised coexpression analysis can be costly and limited to metabolic pathways expressed in the recommended minimum of 15-20 expression conditions required for unsupervised analysis (48). Alternatively, the *CASSIS* algorithm predicts clusters from genomic data by inferring shared intergenic motifs within loci (44,47,49). Function-agnostic approaches can infer noncanonical gene cluster classes *de novo*, although *CASSIS* is primarily implemented in conjunction with function-centric detection to infer cluster boundaries (50).

Another function-agnostic approach identifies gene clusters in genomic data by inferring selection on gene order. Gene clusters comprise cooperative genes that are tied to discrete phenotypes, and the clustered state is maintained by selection (51–53). Gene clusters are evidence of selection because it is improbable that multiple distantly related genes convergently co-locate in diverse lineages and endure microsynteny decay over time (29,51). One proxy that identifies selection on gene order is unexpectedly shared microsynteny (synteny) (29), which detects selection for gene colocalization by identifying combinations of gene families that co-occur with one another in an unexpectedly large phylogenetic distribution. Unexpectedly shared synteny is the basis of the algorithms *CO-OCCUR* and *EvolClust* (28,54) and can identify gene clusters with large or sparse phylogenetic distributions (11,23,55). However, broadly distributed gene clusters are sometimes embedded in larger regions that also meet unexpectedness thresholds because strong linkage disequilibrium within the cluster may extend to the surrounding regions (56). Additionally, if thresholds of unexpectedness are too low, then shared synteny loci that are not gene clusters, such as regions surrounding the centromere, will erroneously be reported as gene clusters (57). To separate gene clusters from loci that are not clusters, unexpectedly shared synteny algorithms have had to sacrifice either their scope of predicted cluster categories, or restrict the sizes of clusters they can predict and/or their capacity to infer recently evolved clusters (55). These limitations are in part why these algorithms have not been as widely implemented as *antiSMASH*.

We created *CLOCI* (Co-occurrence Locus and Orthologous Cluster Identifier) to test two hypotheses: 1) relaxing thresholds for unexpectedly shared synteny will improve cluster recall; 2) measuring additional proxies of selection for coordinated gene evolution will increase sensitivity and accuracy. *CLOCI* lowers unexpectedness thresholds by accurately defining shared synteny locus boundaries and phylogenetic distributions. *CLOCI* enables separating gene cluster families from unexpectedly shared synteny loci that are not gene clusters using three additional proxies for selection for coordinated gene evolution. This approach yields the greatest recovery and categorical diversity of gene clusters when compared to *antiSMASH* and *EvolClust*. The *CLOCI* detection model has the additional advantage of accurate gene cluster boundary inference, and predicts both novel putative clusters and noncanonical biosynthetic gene clusters.

## MATERIAL AND METHODS

*CLOCI* infers gene cluster families (GCFs) by identifying homologous, unexpectedly shared synteny loci and enriches gene clusters using a model of phylogenetic and alignment-based proxies of gene coinheritance and coevolution. The *CLOCI* algorithm first extracts loci with more shared microsynteny than expected (Figure 1a-1i). It then classifies homologous locus groups (HLGs) by adapting graph clustering methodology from orthologous gene group (orthogroup) classification (58,59) (Figure 1j). *CLOCI* enriches HLGs for GCFs using proxies of coordinated gene evolution, including: 1) the unexpectedness of the phylogenetic distribution of an HLG; 2) commitment of genes to the HLG locus; 3) conservative amino acid substitution bias within an HLG; and 4) the sparseness of the HLG distribution across the species phylogeny (Figure 1k). We implemented *CLOCI* across a dataset of 2,247 fungal genomes (Supplementary Table 1) and benchmarked metabolic gene cluster recovery against the function-centric software, *antiSMASH*, and the unexpectedly shared synteny algorithm, *EvolClust,* using a dataset of 57 biosynthetic and 11 non-biosynthetic reference clusters that include catabolic and nutrient assimilation clusters. We benchmark the accuracy of *CLOCI* boundary detection against the same software by referencing a dataset of 33 well-characterized gene cluster boundaries.

**Figure 1.**
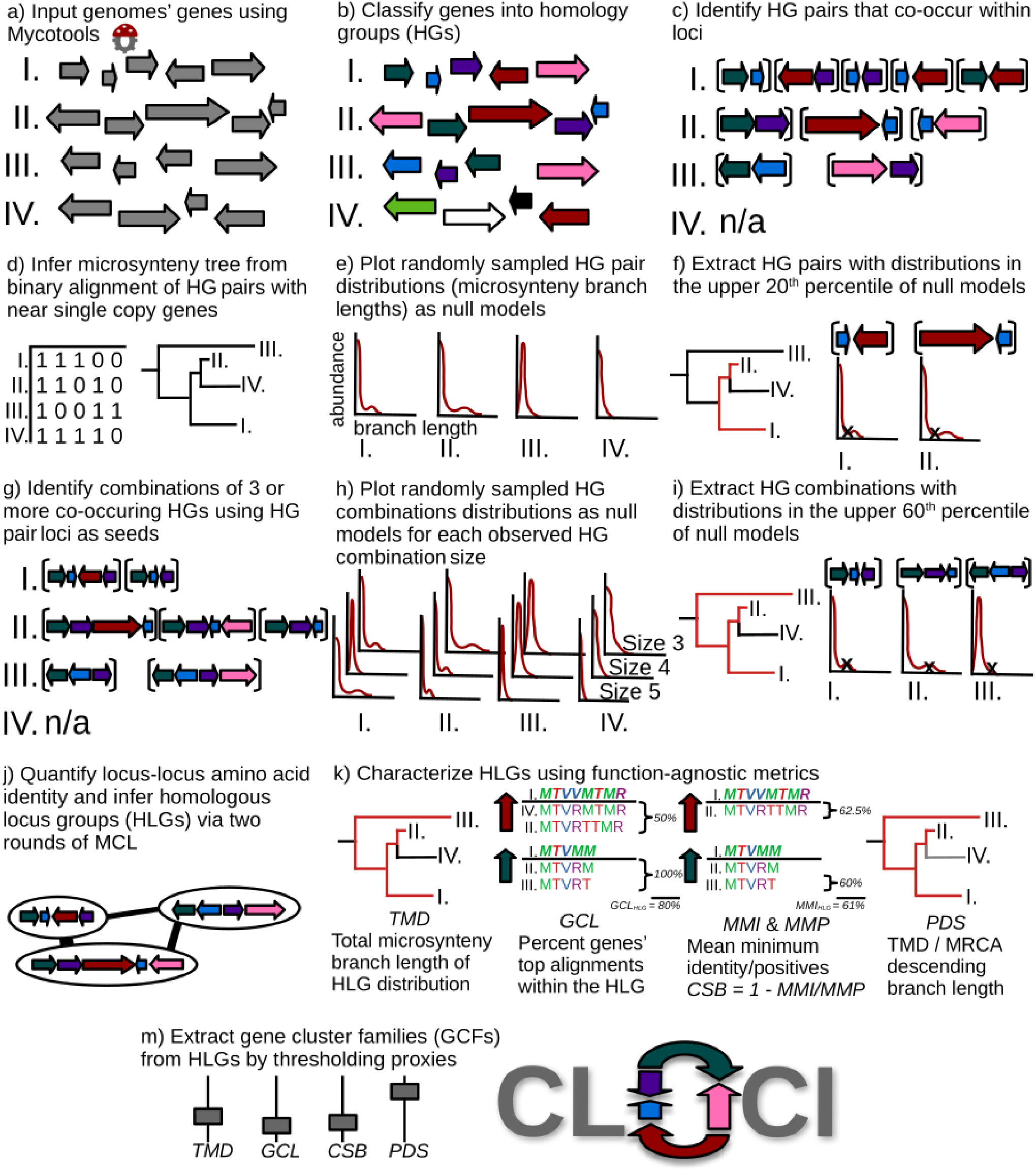
*CLOCI* pipeline depicting cluster detection, including a hypothetical cluster found in genomes *I, II, III*, but not genome *IV*. GCL and MMI in k. are truncated examples using two shared genes from the full HLG.

### Seeding cluster detection by identifying unexpectedly shared microsynteny

Significantly co-occurring pairs of homologous genes (HG) are the seeds for *CLOCI* gene cluster detection. HGs are inferred from a reference MycotoolsDB database (gitlab.com/xonq/mycotools) of complete publicly available genomes using *mmseqs2 cluster* (60) (Figure 1a-1b). The contigs of each genome are then represented as a vector of HGs. All pairs of HGs that co-occur within a five-gene sliding window make up the set of co-occurring HG pairs in a genome (Figure 1c).

In order to quantify the distribution of each HG pair, *CLOCI* constructs a microsynteny tree of the inputted genomes. The microsynteny tree depicts the gene order similarity among genomes, and thus allows us to account for shared evolution when quantifying HG pair distribution. To construct a microsynteny tree, *CLOCI* builds a presence-absence matrix of all HG pairs that include an HG that is a near single-copy ortholog (Figure 1d). We represent microsynteny decay using gene neighborhoods with near single-copy orthologs because these genes are nearly universally conserved and exist in low copy numbers, which limits erroneous identification of orthologous loci. The set of near single-copy ortholog HGs can be determined by BUSCO (61), UFCG (62), or another external method. We used nine UFCG orthologs and five orthologs from a previous study (63) (Supplementary Table 2). Alternatively, *CLOCI* will attempt to determine a representative set of near single-copy HGs present in all genomes in the lowest median and mean count per genome.

The presence-absence matrix is used to reconstruct a maximum likelihood microsynteny tree (64) via *db2microsyntree.py* from the Mycotools software suite (gitlab.com/xonq/mycotools). The microsynteny tree is built using IQ-TREE 2 and a free-rate, general time reversible evolutionary model with ascertainment bias correction (65–67) (Figure 1d). To prevent topological discrepancies with accepted species topology (62,68), the microsynteny topology can be constrained with a species tree or in the absence of an external constraint a consensus tree constructed from 1000 ultrafast bootstrap replicates is generated. We constrained the microsynteny topology with a phylogenomic tree constructed from the same near single-copy orthologs used to detect gene neighborhoods for microsynteny tree construction (Supplementary Table 3). The tree is then rooted on specified genomes, and the distribution of each HG pair is calculated as the total microsynteny branch distance (TMD) between the genomes where it is found (Equation 1).

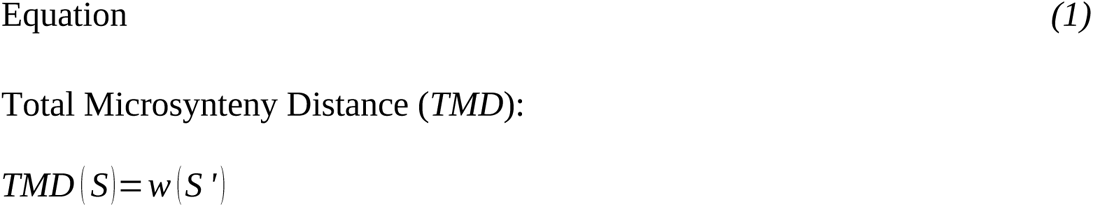

Where *MT = (V, E)* is a rooted microsynteny tree with vertices *V,* edges *E,* and a positive weight *w(e)* (branch length) assigned to each edge, *e*. *L(MT)* denote the set of leaves (genomes) of *MT.* For any set of genomes *S*={*s*_0_, *s*_1_… *, s_n_*}*⊆L* (*MT*) that contain an HG pair, HG combination, HLG, or GCF, *S’* denotes the smallest subtree of *MT* containing *S. w(S’)* is the sum of the weights of the edges of *S’*.

To identify unexpectedly distributed HG pairs, *CLOCI* creates a null distribution of background TMD from randomly sampled HG pairs (Figure 1e). Different lineages have different rates of microsynteny decay, so *CLOCI* builds local null models for specific lineages or taxonomic ranks. By default, *CLOCI* builds null models for each genus because microsynteny decay often does not significantly vary within genera (69), and it strikes a balance with computational throughput. *CLOCI* accounts for species with overrepresented genome samples during null model construction by randomly choosing a species prior to randomly sampling from a genome. The upper 20^th^ percentile from each null distribution is used to define the TMD value above which an HG pair is considered unexpectedly shared (Figure 1f). We selected the 20^th^ percentile to decrease downstream pairwise comparisons while presumably identifying most HG pairs that represent gene clusters.

HG pairs are used as seeds for inferring higher-order HG overlap (Figure 1g). Combinations of three or more co-occurring HGs (HG combination) are identified by pairwise comparison of all loci that correspond to an HG pair. Null distributions are generated for each observed HG combination size, up to the sliding window, as described for HG pairs (Figure 1h). Unexpectedly syntenic HG combinations are extracted from the upper 60^th^ percentile of null TMDs (Figure 1i). We chose the 60th percentile as the threshold below what is necessary to detect the recently evolved psilocybin reference cluster (41,43).

### Identifying locus boundaries by assembling shared synteny loci

Shared synteny locus boundaries are predicted by first extracting loci from unexpectedly shared HG combinations (Figure 1j). All loci that correspond to unexpectedly shared HG combinations are extracted. Then any overlapping genes between these loci are added to the locus that contains the HG combination with the greatest TMD. Single gene loci generated by the merging process are fused with an adjacent locus with the most similar phylogenetic distribution (Equation 2) if it exceeds a 25% minimum phylogenetic similarity. If a single gene is not merged, then it is aggregated with surrounding singletons, or discarded if no surrounding singletons exist. Spurious loci generated from singleton merging are discarded downstream if they lack homology to other loci.

*CLOCI* finalizes shared synteny locus boundary inference by identifying groups of similar domains of shared microsynteny and merging adjacent loci with homologous domains. To infer groups of similar domains, *CLOCI* implements Markov clustering (MCL) (70) on an adjacency graph of locus-locus similarity. *CLOCI* quantifies the pairwise similarity of all loci following their extraction. We define similarity between two loci as the average amino acid identity of their shared HGs scaled by the overlap coefficient of the shared HG combinations (Equation 3). Compared loci must have at least two shared HGs, and if a locus contains multiple gene sequences from a particular HG then the highest identity comparison is incorporated into the average. Locus-locus similarity scores greater than 35% are represented in an adjacency graph, and initial domains are inferred via this graph with inflation value set to 1.1. *CLOCI* merges extracted loci that are within two genes of one another and the loci belong to the same domain.

### Grouping locus homologs by iterative graph clustering

To identify groups of similar loci and families of related gene clusters, by extension, *CLOCI* groups the finalized extracted loci into homologous locus groups (HLGs). HLGs are inferred using the same locus-locus similarity and graph clustering algorithm as used for detecting initial domains, though in this round *CLOCI* implements the Sørensen-Dice similarity as the scaling coefficient (Equation 3) to penalize against missing HGs in the similarity calculation. HLGs are inferred from the resulting locus-locus adjacency graph with a default MCL inflation value of 1.3.

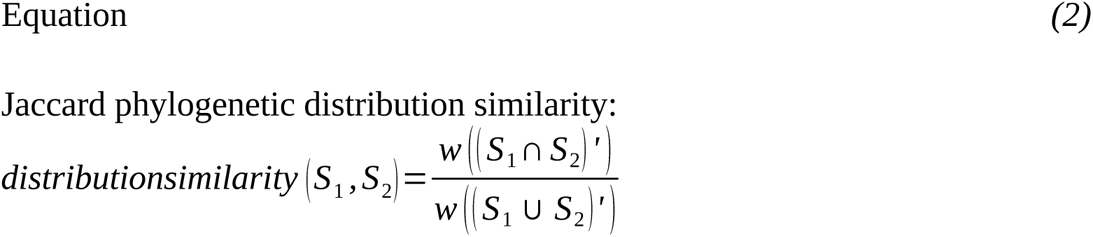

_For any two sets of genomes_ 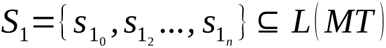 _and_ 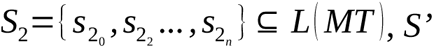 _denotes_ the smallest subtree of *MT* containing *S.* Let *w(S’)* be the sum of the weights of the edges of *S’* (TMD).

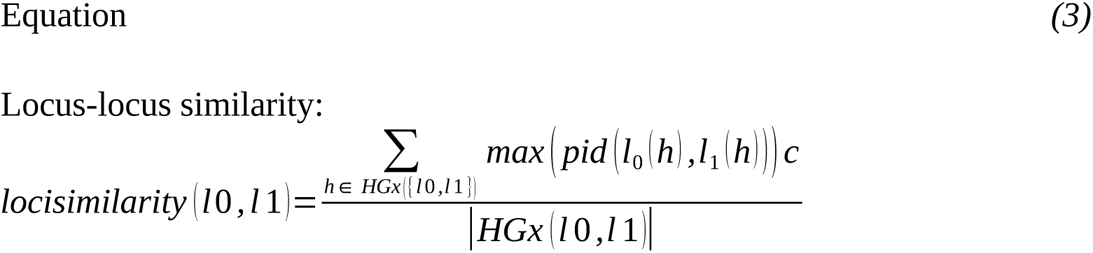

Where *l0* and *l1* indicate the loci compared; *h* indicates an HG; *HGx* represents the HG combination of a locus/loci such that *HGx* (*l* 0)={*h*_0_ *, h*_1_,…*h_n_* } and *HGx* ({*l* 0 *,l* 1})=*HGx* (*l* 0) *∩ HGx* (*l* 1)*; l0(h)* denotes the gene(s) within locus *0* corresponding to *h;* and *pid* is the percent alignment identity between two genes. *c* depicts a similarity coefficient of each set of *HGxs,* such as the overlap coefficient, Sørensen-Dice coefficient, or Jaccard Index.

*CLOCI* next attempts to repair incomplete or fragmented loci within inferred HLGs. First, *CLOCI* attempts to complete partial cluster loci by reference to related loci. Each locus within each HLG is extended to include any HG within one gene up/downstream of the locus that is shared across the final HLG, unless the flanking gene is already part of a different HLG. The HG is later removed if it has less than 35% amino acid identity with all homologous sequences in the HLG. To identify loci that are fragmented across the genome assembly, all contigs are searched for shared HG pairs that are missing from a given locus. *CLOCI* only searches for two gene combinations because combinations greater than three are already retrieved by identifying unexpectedly syntenic HG combinations.

### Characterizing homologous locus groups according to proxies of coordinated gene evolution

Following HLG circumscription, *CLOCI* characterizes HLGs using proxies of coordinated gene evolution in addition to TMD, including gene commitment to the locus (GCL), locus-locus amino acid similarity, conservative amino acid substitution bias (CSB), and the Phylogenetic Distribution Sparsity (PDS) of HLGs (Figure 1k). To visualize the distribution of HLG proxies and relative contribution of each HLG proxy to this distribution, we reduced the proxy dimensionality using principal component ordination implemented in the *scikit-learn* package (71). We tested for autocorrelation between proxies using ordinary least squares linear regression.

#### Gene Commitment to the Locus

Gene Commitment to the Locus (GCL) is a measure of the extent to which an HG has remained in an HLG through time. This is a proxy for the coordinated inheritance of genes in the HLG. The GCL for an HLG is the weighted average of individual HG commitment scores for each HG found in at least two homologous loci of the HLG (Equation 4). The HG commitment score is equal to the percent of alignment hits that are sequences within the HLG after discarding paralogs outside the HLG locus in genomes that have it. Sequences are aligned against the entire HG using *DIAMOND BLASTp.* Hits are determined by sorting the alignments from highest to lowest identity, and retaining sequences up until all homologous sequences within the HLG loci are recovered. Only the maximum scoring sequence is considered if a genome has multiple sequences in the same HG within an HLG.

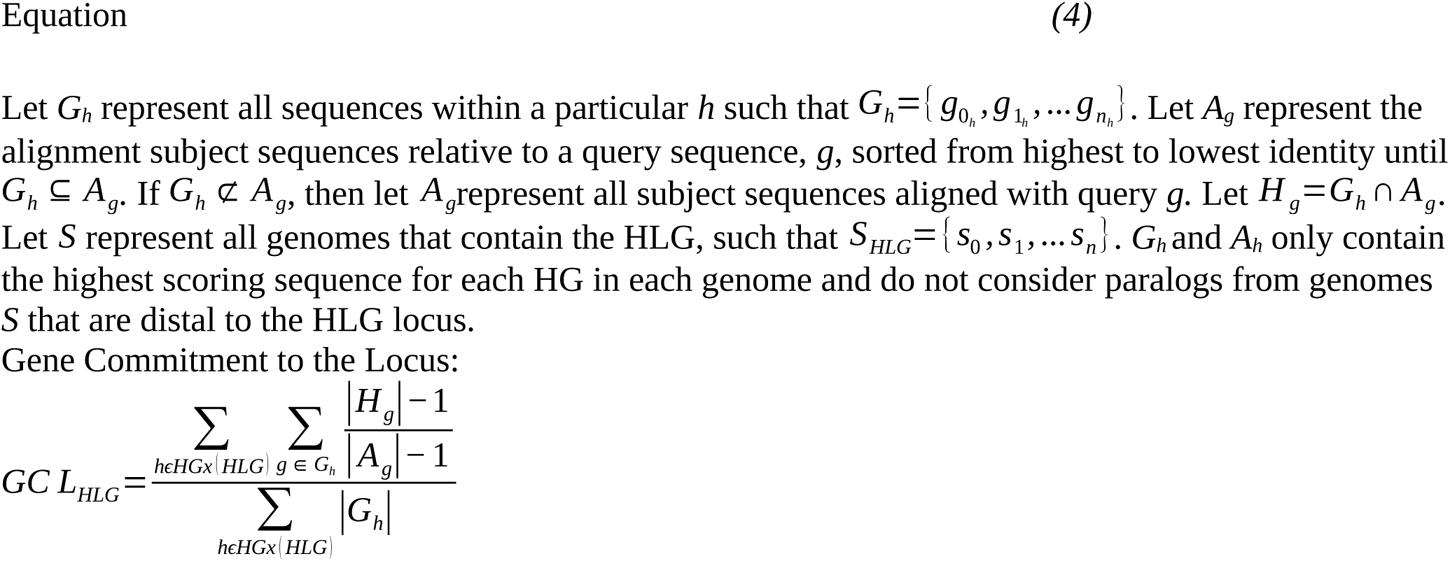

Where *HLG* is a set of homologous loci, such that *HLG*={*l* 0 *, l* 1,… ln}.

#### Conservative Substitution Bias

Conservative Substitution Bias (CSB) is a proxy for the action of purifying selection on the protein structure that is more computationally tractable than dN/dS. CSB measures the ratio of positive amino acid substitutions to identical amino acids. CSB for an HLG is calculated by dividing the mean minimum observed percent amino acid identity (MMI) by the mean minimum observed percent conservative alignment positions (positives, MMP) across all pairwise comparisons of sequences in an HG. Minimum values approximate the maximum distance between sequences in the HLG. To calculate the minimum identities and positives, all sequences in HGs in two or more homologous loci in the HLG are aligned against the entire HG via *DIAMOND BLASTp*. The minimum observed positive score for all pairwise alignments is divided by the minimum identity score. The minimum identity and positive scores are set to 0 for alignments that are missing any sequences in the HLG. The HLG CSB is then derived from the ratio of HG minimum identities to minimum positives scaled by the proportion of considered genes in each HG (Equation 5).

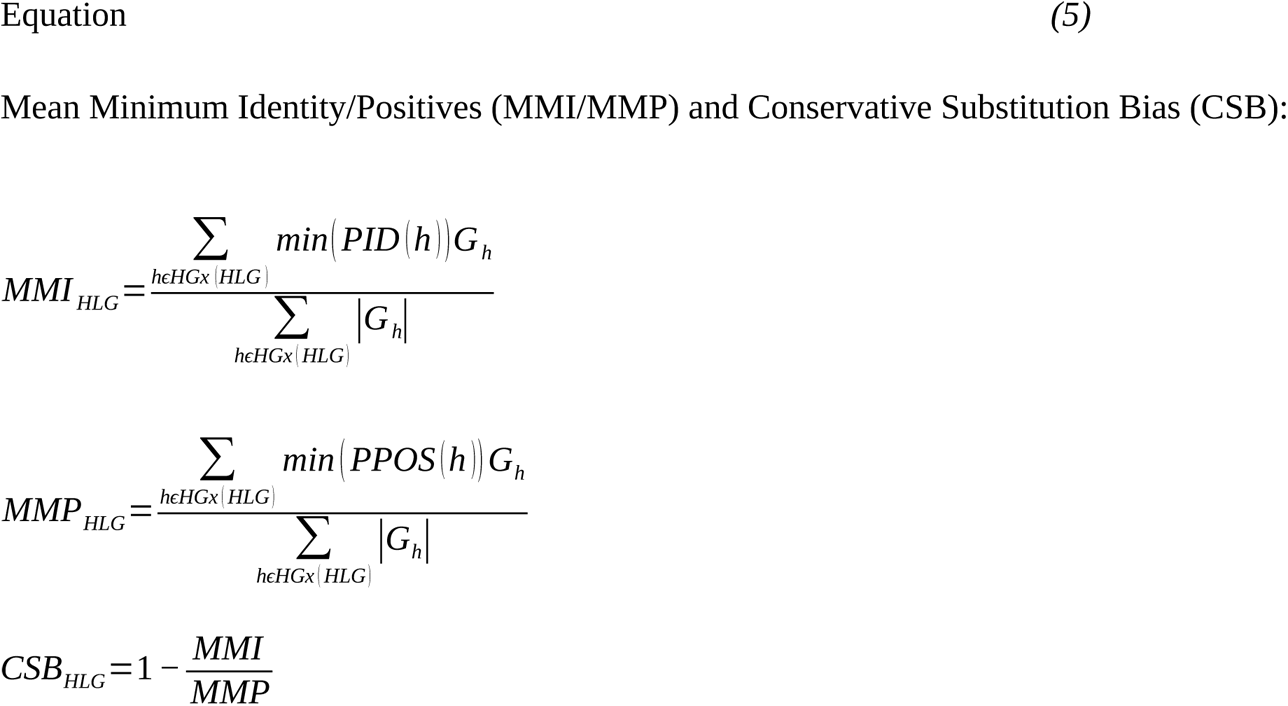

Where 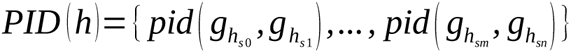 such that *PID* (*h*)represents the set of all intergenome percent amino acid identities for a particular *h* and *ppos* (*g*_0_ *, g*_1_) indicates the percent amino acid positives between g_0_ and g_1_, and 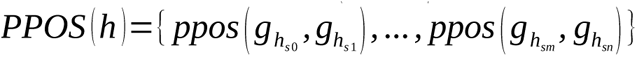.

#### Phylogenetic Distribution Sparsity

Phylogenetic Distribution Sparsity (PDS) measures the prevalence of taxa with a particular HLG in its overall phylogenetic range. PDS is a proxy for the extent of horizontal cluster transfer and/or coordinated cluster gene loss, which are elevated in some categories of gene clusters (25,43,51,55,72,73). *CLOCI* quantifies PDS for each HLG as the TMD of the HLG divided by the TMD of the most recent common ancestor of the genomes that have the HLG (Equation 6). Thus, PDS quantifies the percent of branch length descended from the most recent common ancestor TMD that contains the HLG.

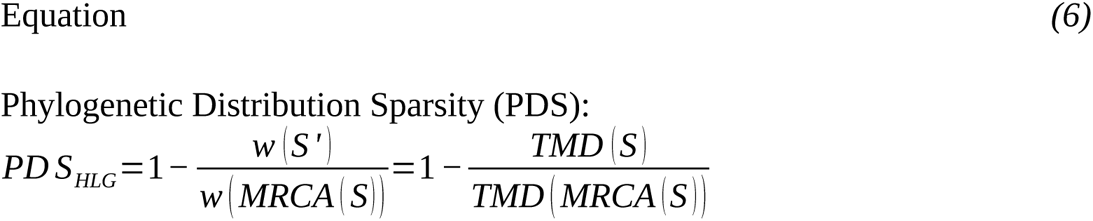

where *MRCA(S)* is the smallest subtree of *T* containing all tips descended from the most recent common ancestral node of *S.* Let *w(MRCA(S))* represent the sum of the weights of the edges of *MRCA(S)*.

### Implementing *CLOCI* using a dataset of publicly available fungal genomes

We implemented *CLOCI* across a database of 2,247 fungal genomes across the kingdom using default *CLOCI* parameters (Supplementary Figure 1, Supplementary Table 1) using Mycotools. We additionally constrained the microsynteny tree topology to 14 near single-copy orthologs from Spatafora et al. 2013 and Universal Fungal Core Genes (62) and rooted on *Rozella* spp (Supplementary Table 2). We created GCF networks locus-locus similarity networks using the *hlg2hlg_net.py* tool packaged with *CLOCI.* All computation was performed using 26 Quad Core Intel Xeon 6148 Skylake processors with access to approximately 1 terabyte RAM.

### Benchmarking *CLOCI* recovery and boundary accuracy against *antiSMASH* and *EvolClust*

We benchmarked detection power with reference to a dataset of known clusters (Supplementary Table 4) and the precision and accuracy of cluster boundaries with reference to an independent dataset that primarily contains well-characterized *Aspergillus* spp. gene clusters (Supplementary Table 5). We cannot directly test the false discovery rate and algorithm precision because gene clusters are not exhaustively sampled in any analyzed genomes. We compared *CLOCI* reference cluster recovery and cluster boundary precision with those of *antiSMASH* (v6.1.1) and *EvolClust* (1,74). We implemented *antiSMASH* on the genomes containing known clusters and queried the *EvolClustDB* (accessed 2023, April 4). If the *EvolClustDB* genome was from a different strain than the query cluster, we first verified that the cluster was present by querying an *EvolClustDB* genome of the same species using *BLASTp*. We evaluated the recovery of 57 unique biosynthetic clusters and 11 non-biosynthetic clusters. Biosynthetic clusters were manually selected from MiBIG to maximize diversity of known biosynthetic classes. We included the psilocybin and biotin clusters to increase noncanonical biosynthetic cluster representation (19,43,75). The non-biosynthetic reference clusters were compiled from an extensive search of available literature (23). We tested recovery by aligning a core biosynthetic gene or gene near the center of each reference cluster to the same genome as the reference (Supplementary Table 1, Supplementary Data). Cluster loci were deemed recovered if the top *BLASTp* hit to the reference gene had a minimum percent identity 90%. If the reference genome was not present in our dataset, we confirmed the presence of the reference cluster in a genome of a species known to have the cluster by identifying two top *BLASTp/tBLASTn* hits within the same locus with greater than 60% identity to two reference cluster genes (Supplementary Data). Four clusters were removed from the *antiSMASH* recovery analysis because the genome annotation format was incompatible with *antiSMASH*. 19 clusters were removed from the *EvolClust* recovery calculation because the genome containing the query cluster was not in *EvolClustDB*.

We assessed cluster boundary accuracy referencing a separate dataset of 33 gene clusters, 25 of which are derived from the *CASSIS Aspergillus* boundary accuracy dataset (47), because most clusters do not have well characterized boundaries (Supplementary Table 5). We additionally included psilocybin (41) and biotin (19) noncanonical biosynthetic gene clusters, three non-biosynthetic gene clusters (16,25,76), the *Penicillium chrysogenum* penicillin cluster (11), and two ergot alkaloid biosynthetic gene clusters (31) to broaden the scope of gene cluster categories in boundary detection. We assessed accuracy by the percent missing and percent extraneous genes per cluster. Clusters were submitted to boundary evaluation if at least one gene sequence within the boundaries was identified using *BLASTp*. All reported loci that contain at least one of the genes within the reference cluster boundaries are considered in boundary accuracy.

### Evolutionary analysis of nitrate assimilation gene clusters

We examined the evolution of nitrate assimilation gene clusters by constructing gene phylogenies (77) for cluster genes using the Cluster Reconstruction and Phylogenetic Analysis pipeline packaged in Mycotools (gitlab.com/xonq/mycotools) with default parameters (Supplementary Table 6). We extracted nodes with high support (> 0.99 fasttree bootstrap) and generated subsequent phylogenies by aligning with *mafft –auto* v7.487 (78), trimming using *ClipKIT* v1.3.0 (79), and constructing phylogenies using IQ-TREE v2.2.0.3 with 1000 ultrafast bootstrap replicates (65,67). To test evaluate the alternative hypothesis of vertical evolution compared to horizontal transfer, we compared the likelihood of a Mucoromycota monophyletic constraint using an approximately unbiased test (80) with 10,000 boostrap replicates implemented in IQ-TREE v2.2.0.3 (67) (Supplementary Table 7).

### Filtering gene cluster families from homologous locus groups according to proxies of coordinated gene evolution

Gene cluster families (GCFs) are filtered from HLGs by setting thresholds for the minimum proxy values (Figure 1l). In order to evaluate the capacity for coordinated gene evolution proxies to enrich for metabolic gene clusters, we analyzed the effect of variable coordinated gene evolution proxy thresholds on the proportion of metabolic process (GO:0008152) and secondary metabolic process (GO:0019748) gene ontology (GO) terms in the resulting GCFs. To obtain GO terms, we first queried the genes against the Pfam database using *hmmsearch* v3.3.2 with 50% minimum alignment coverage and 0.001 minimum e-value. Each gene was annotated with its lowest e-value Pfam hit and GO terms were assigned referencing the pfam2go database (current.geneontology.org/ontology/external2go/pfam2go). We incrementally adjusted TMD, GCL, CSB, and PDS thresholds by 0.2 from zero to one and quantified the proportion of retained genes annotated as metabolic process or secondary metabolic process. The TMD calculation in this filtering model acts on the distribution of an HLG rather than individual HG pairs/combinations as it does in unexpectedly shared microsynteny detection.

## RESULTS

*CLOCI* recovers functionally diverse families of gene clusters. We recovered all reference gene cluster categories, including both canonical biosynthetic (aflatoxin) and noncanonical biosynthetic (psilocybin), catabolic processes (quinic acid, galactose, and proline degradation), and nutrient assimilation (nitrate assimilation) clusters. These gene clusters were recovered from an output of 119,104 homologous locus groups (HLGs) that comprise 3,213,162 unexpectedly shared synteny locus domains with an average size of 4.82 genes per domain (61.2% of overall genes), median size 4 genes, maximum size 129 genes, and minimum size of 2 genes. There are 27.0 locus domains per HLG, and most reference boundary clusters are represented by a single locus domain, though we recovered some clusters as two or three domains. The *CLOCI* gene cluster detection algorithm has higher cluster recovery than the function-centric algorithm, *antiSMASH*, and the unexpected shared synteny algorithm, *EvolClust*. *CLOCI* also better approximates cluster boundaries when considering all shared synteny locus domains in reference clusters. Filtering using the minimum observed proxy values for reference clusters removes 39,429 HLGs (33.1%). This filtered dataset comprises 79,675 gene cluster families (GCFs), with 35.4 cluster domains per GCF, mean size 4.93 genes per cluster domain (54.9% of overall genes), median size 4 genes, maximum size 129 genes, and minimum size 2 genes. Raising threshold proxy values variably enriches for genes with metabolic process and specialized metabolite process GO terms in the final set of GCFs.

### *CLOCI* gene cluster families recapitulate known gene cluster distributions

To assess gene cluster family (GCF) completeness, we compared the reported phylogenetic distribution of gene clusters to their respective *CLOCI* GCFs. We recovered known gene cluster distribution in the main categories of gene clusters. We recovered the ergot alkaloid biosynthetic GCF in all Onygenales, Eurotiales, Xylariales, Hypocreales, and Helotiales genomes that we previously identified through phylogenetic analysis (71). We identified all known homologs of the noncanonical biosynthesis gene cluster for psilocybin production in our dataset in a single GCF (Fig. 3a). Additionally, both parts of the *Pa. cyanescens* psilocybin cluster, which is distributed across two contigs due to a fragmented genome assembly, were recovered in this GCF. We also identified unreported homologs of the clusters that produce the immunosuppressant nonribosomal peptide, cyclosporin, in *Dactylonectria estremocensis* (Supplementary Data), and the fungicidal strobilurin polyketides in *Mycena* spp. (Figure 4c).

Multiple nitrate assimilation GCFs were identified from diverged taxa (25). We recovered at least five GCFs that contain nitrate transporter, nitrate reductase, and nitrite reductase genes associated with nitrate assimilation. These GCFs span three phyla and the clusters are often found directly adjacent to unexpectedly shared synteny regions. GCF #0 consists entirely of Ascomycota clusters, including the characterized *Aspergillus nidulans* cluster (18). GCF #1 comprises 81.3% Basidiomycota clusters and includes sequences implicated in horizontal transfer between ancestral Ustilagomycotina (Basidiomycota) and Hypocreales (Ascomycota) species (66,70). GCF #2 contains the five-gene Saccharomycotina (Ascomycota) yeast nitrate assimilation gene cluster reported from *Ogataea polymorpha* (81), as well as unreported homologous clusters in *Bifiguratus adelaidae* (Mucoromycota). GCF #3 primarily contains *Pseudogymnoascus* spp. (Ascomycota) (82). Additionally, an unreported putative nitrate assimilation GCF #4 was identified Basidiomycota and Mucoromycota (Figure 4f), consisting of a nitrate transporter, nitrate reductase, and nitrite reductase. All nitrate assimilation GCFs are connected in the locus-locus similarity network, indicating shared homology (Figure 4f). Phylogenetic reconstruction of the nitrate transporter, nitrate reductase, and nitrite reductase genes in GCF #4 using *crap.py* (gitlab.com/xonq/mycotools) reveals that ectomycorrhizal *Amanita* spp. (Basidiomycota) are nested within a clade of Mucoromycota spp. with 100% IQ-TREE ultrafast bootstrap support for all three genes (67). Constrained trees that force a Mucoromycota monophyly were rejected via approximately unbiased testing (Supplementary Table 7). *Amanita muscaria* spp. have close homologs of these genes but lack an intact cluster and saprobic *A. thiersii* and *A. inopinata* both lack homologs of the genes (83).

### Benchmarking *CLOCI* against *antiSMASH* and *EvolClust*

We benchmarked *CLOCI* cluster recovery against *antiSMASH* and *EvolClust* referencing a dataset of 11 non-biosynthetic gene clusters from literature and 57 biosynthetic clusters, including two noncanonical biosynthetic clusters (Supplementary Table 4). *CLOCI* recovers 91.2% and 100% of the biosynthetic and non-biosynthetic dataset respectively (Figure 2, Supplementary Data). In comparison, *antiSMASH* recovers 76.1% and 0% of the biosynthetic and non-biosynthetic datasets respectively, whereas *EvolClustDB* recovers 48.7% and 60% respectively. Additionally, *CLOCI* detects the biotin and psilocybin noncanonical biosynthetic clusters. Of the missing clusters, *CLOCI* did not detect the zearalenone, mycophenolic acid, cephalosporin, or the ent-kauren-16alpha-ol clusters.

**Figure 2.**
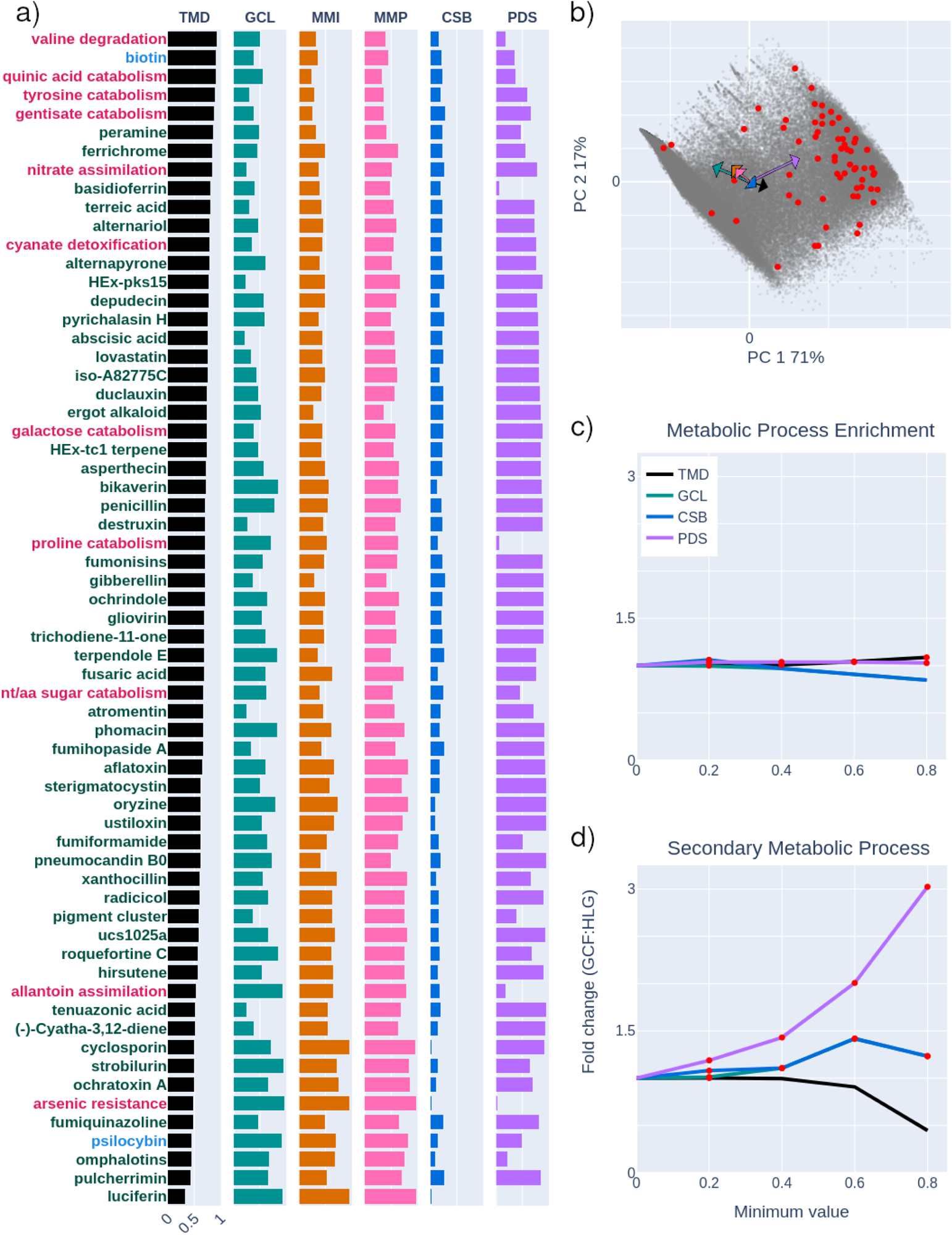
Reference gene cluster families (GCFs) are characterized by and extracted from shared synteny homologous locus groups (HLGs) using proxies of coordinated gene evolution: a) Bar graphs of proxies of coordinated gene evolution for 63 recovered clusters. Nitrate assimilation is represented by the GCF containing the *Ustilago bromivora* cluster. TMD = Total Microsynteny Distance, GCL = Gene Commitment to the Locus, MMI/MMP = Mean Minimum Identity/Positives of amino acids, CSB = Conservative Substitution Bias, and PDS = Phylogenetic Distribution Sparsity. Red font indicates non-biosynthetic clusters, green canonical biosynthetic, and blue noncanonical biosynthetic. b) Principal component analysis of HLGs highlighting HLGs with reference clusters in red. Loading (feature) vectors correspond to the proxy with the same color as bars in *a)*; Enrichment of c) metabolic processes (GO:00008152) and d) secondary metabolic processes (GO:00019748) in GCFs filtered from HLGs with greater than a minimum specified proxy threshold relative to initial HLGs. Significantly enriched data points are in red.

We assessed cluster boundary inference by referencing an independent dataset of well-characterized cluster boundaries (Supplementary Table 5). We determined that *CLOCI* has the lowest mean (0.55) and median (0) missing genes and lowest mean (0.90) and median (0) extraneous genes per recovered cluster (Figure 2b). 65.6% of reference clusters were predicted as a single domain by *CLOCI*, 27.6% were reported as two, and one cluster was reported as three domains. For example, aflatoxin was reported as a single domain, whereas sterigmatocystin was reported as two domains, one of which is circumscribed in the same GCF as aflatoxin. The pseurotin and fumagillin clusters are separated by a single gene and *CLOCI* reports these clusters as two discrete loci that comprise three and one domains respectively. All clusters were recovered as a single domain by *antiSMASH* and *antiSMASH* with *CASSIS* enabled, and pseurotin and fumagillin clusters were reported as a united locus. *AntiSMASH* recovers the aflatoxin cluster in the boundary dataset, but did not recover the query gene for the recovery dataset because the recovery dataset cluster references a fragmented assembly. The aflatoxin and cichorine clusters were recovered as two domains by *EvolClust*.

### Filtering using proxies of coordinate gene evolution reveals novel gene clusters

*CLOCI* filters GCFs from shared synteny HLGs using multiple proxies of selection for coordinated gene evolution (Table 1, Figure 2a-b). Proxies have variable power to filter the dataset when their observed minimum values in the reference dataset are used independently. The minimum Total Microsynteny Distance (TMD) reduces HLGs by 19.0%, Gene Commitment to the Locus (GCL) by 10.8%, and Conservative Substitution Bias (CSB) by 8.89%. Phylogenetic Distribution Sparsity (PDS) is 0% for luciferin. However, overall, PDS tends to be greater than 0% (median 81.3%), and PDS also explains most of the variance in the distribution of both principal components following two dimensional ordination of HLG proxies.

**Table 1:**
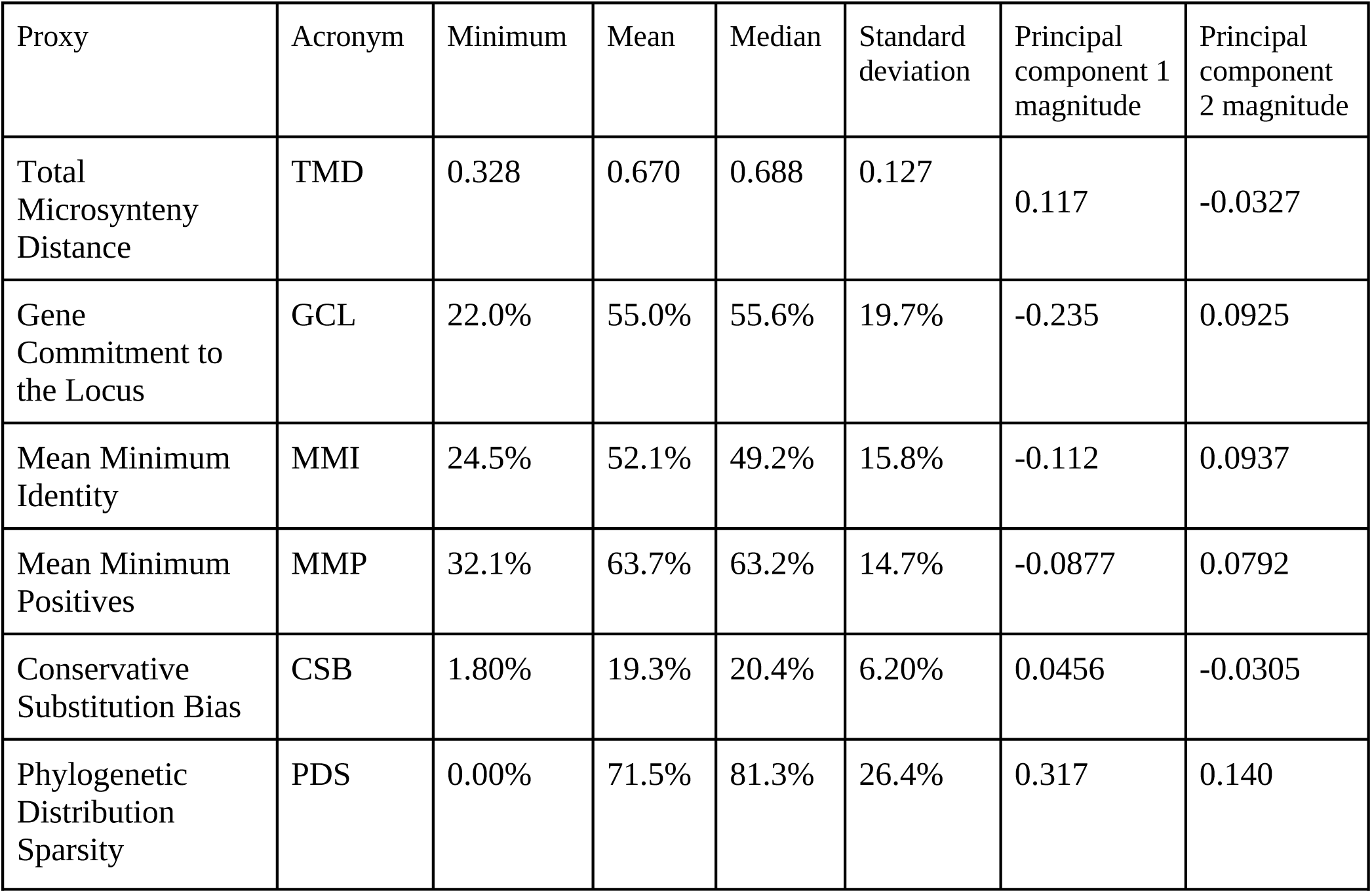
Values of coordinated gene evolution proxies for homologous locus groups (HLGs) that contain known gene clusters from the recovery dataset.

Reference specialized metabolic clusters have the lowest TMDs, while clusters that encode vitamin biosynthesis, catabolic, and cellular protective functions have the highest (Figure 2a). The four clusters with the smallest TMD are the secondary/specialized metabolite clusters luciferin, pulchirmerrin, omphalotins, and psilocybin. The top four TMD clusters encode valine catabolism, biotin biosynthesis, quinic acid catabolism, and tyrosine degradation/pyomelanin biosynthesis. Biotin is an essential (19), while the quinic acid cluster catabolizes a carbon source (76,84), the tyrosine degradation/pyomelanin biosynthesis cluster degrade phenolics and produce melanins that may protect fungal cells from the environment (17,85).

TMD is weakly to strongly linearly correlated with other proxies using an arbitrary adjusted R^2^ cutoff of 0.5, while the other proxies tend to lack correlation with one another (Supplementary Figure 1). Log normalized TMD is correlated with GCL (adjusted R^2^ = 0.606), PDS (adjusted R^2^ = 0.542), and CSB (adjusted R^2^ = 0.833). There is no evidence that GCL is linearly correlated with PDS (adjusted R^2^ = 0.127) or CSB (adjusted R^2^ = 0.446). PDS may be weakly correlated with CSB (adjusted R^2^ = 0.509).

Filtering using minimum threshold values for proxies of selection for coordinate gene evolution differentially affects enrichment of metabolic process and secondary metabolic process gene ontology (GO) terms. We determined that metabolic process genes are significantly and increasingly enriched as the TMD threshold is raised (Figure 2c). Conversely, increasing TMD reduces secondary metabolic process genes (Figure 2d). GCL reduces metabolic process genes across the thresholding range. However, CSB and GCL similarly enrich secondary metabolic process genes from 0.2 to 0.8 with a peak fold change at 0.6 for both. On its own, raising the CSB threshold to 0.2 enriches metabolic process genes, though larger thresholds lead to diminishment. PDS enriches both metabolic process and secondary metabolic process genes across the range of thresholds, with a peak metabolic process enrichment at 0.2. At 0.8, PDS generates a 302% increase in the proportion of secondary metabolic process genes.

*CLOCI* putatively identifies an unreported widely shared metabolic cluster and sparsely distributed specialized metabolite cluster. We putatively identified a widely shared iron sequestration gene cluster with an 0.81 TMD that is log normalized relative to all HLG TMDs. This gene cluster contains a “widely conserved” synthetase that produces the Basidiomycete siderophore, basidioferrin (12). The *CLOCI* reported cluster is relatively large, comprising 23 predicted genes in *Psilocybe cyanescens* (Figure 4e). The TMD of the basidioferrin GCF is the 9^th^ greatest when compared to reference clusters and its PDS is near 0 (Figure 3a), which is congruent with the broad distribution of basidioferrin synthetase homologs (12). We also identified a putative non-ribosomal peptide specialized metabolite GCF (# 91,365) in the upper 99 percentile of PDS scores. This putative GCF was recovered in four genomes within three taxonomic classes.

**Figure 3.**
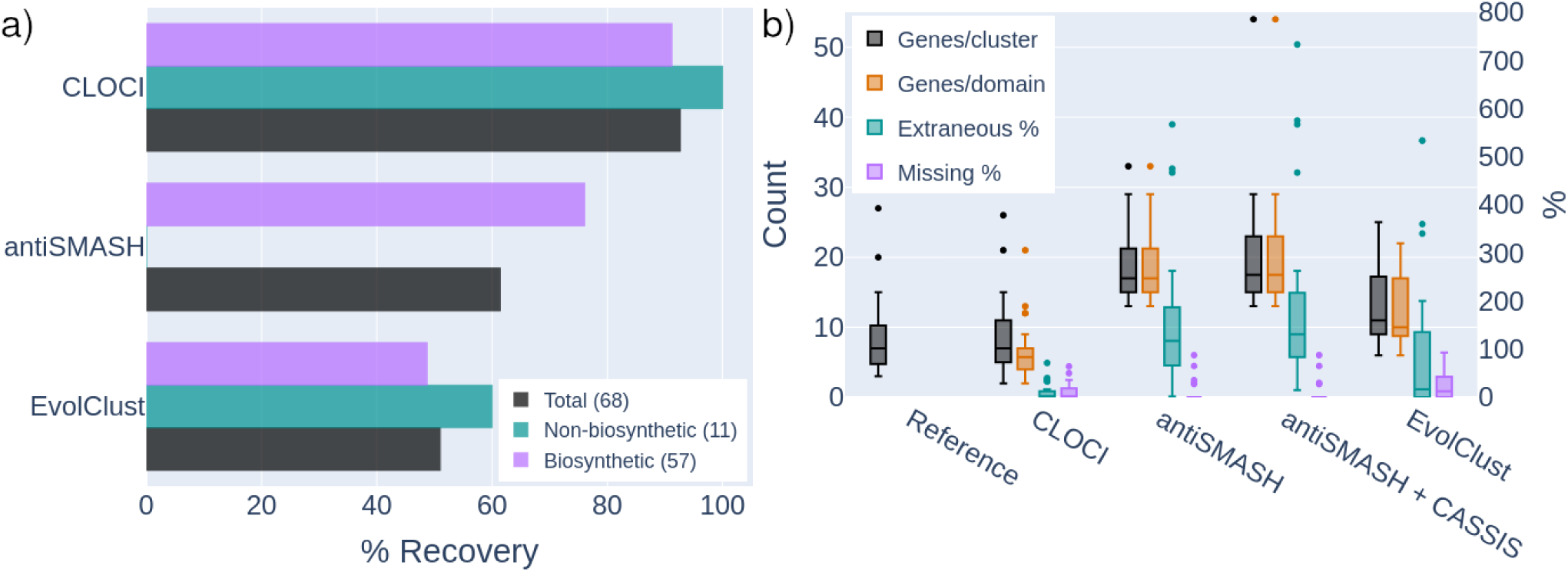
Comparison of cluster recall and cluster boundary accuracy by *CLOCI*, *antiSMASH, CASSIS,* and *EvolClust*. a) Recovery of 57 biosynthetic gene clusters from (38) and literature, and 11 non-biosynthetic gene clusters from (23). b) Boundary detection accuracy and precision referencing an independent dataset of 33 characterized cluster boundaries from (46) and literature. Whiskers depict the 1^st^ and 3^rd^ quartiles.

**Figure 4.**
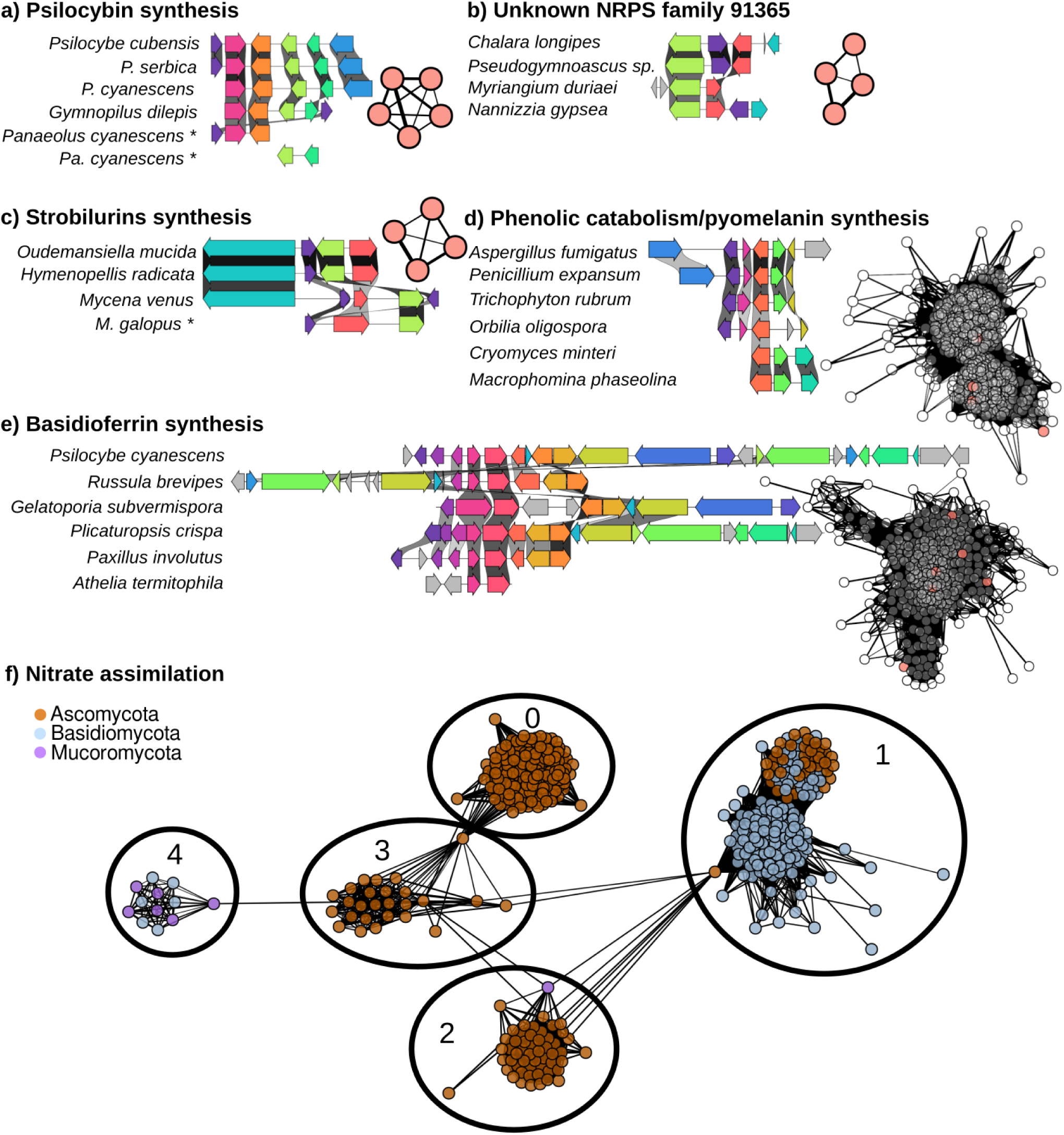
Locus-locus similarity graphs and representative synteny diagrams of *CLOCI* GCFs of multiple gene cluster categories. Nodes represent loci and edge weight/distance depicts log normalized locus-locus similarity (Equation 3). Nodes depicted in synteny diagrams in a-e are salmon-colored and ‘*’ indicates clusters recovered from contig edges. Gene arrows are color coded by homology, and grey arrows indicate no inferred homology. a) The noncanonical psilocybin biosynthetic gene cluster in all reported genomes in the dataset, including across two contigs in *Panaeolus cyanescens*. b) An unreported cluster containing a nonribosomal peptide synthetase found in three different taxonomic classes. c) Strobilurin clusters including unreported homologs in *Mycena*. d) The phenolic catabolism and pyomelanin biosynthesis cluster family. e) A putative basidioferrin gene cluster family that is widely shared across Agaricomycotina (Basidiomycota) genomes. Basidioferrin synteny diagram is centered on the characterized *Gelatoporia subvermispora* basidioferrin type VI nonribosomal peptide synthetase. f) Similarity network of five nitrate assimilation GCFs distributed among three phyla.

## DISCUSSION

### *CLOCI* generalizes gene cluster detection by inferring selection on coordinated gene evolution

Detecting selection for coordinated gene evolution is an effective tool for identifying gene clusters (28,29,86). Unexpectedly shared synteny (USS) detection identifies gene clusters from diverse categories by identifying selection for gene colocalization (29). However, USS detection does not discriminate between gene clusters and loci that have USS for other reasons, so in order to identify true gene clusters, previous USS algorithms limit their capacity to infer recent, small clusters or generally infer cluster categories. *CLOCI* identifies recently evolved clusters while remaining function-agnostic in part by accurately inferring locus boundaries in a size-agnostic framework. Detecting other signatures of selection for coordinated gene evolution can then enrich USS loci for metabolic gene clusters.

The signature of selection for gene colocalization can identify gene clusters from diverse categories, including phenolic and amino acid catabolism, nutrient assimilation, and both noncanonical and canonical biosynthetic gene clusters (28,29). Non-biosynthetic clusters have largely been overlooked by function-centric software because they are primarily designed to detect biosynthetic clusters. Noncanonical biosynthesis has been overlooked because these clusters lack screened core gene models, and often have discrete distributions. USS, employed in *CLOCI* and other methods (28,54,86) is a proxy for selection on gene colocalization, a driver of the assembly and maintenance of gene clusters. Because this general evolutionary signature of gene clusters is not expected to vary based on cluster function, all categories should be detectable by this method, in contrast with function-centric methods (55). Previous USS algorithms, such as *CO-OCCUR,* indeed can infer gene clusters from diverse categories (28,29). However, limitations in USS cluster recovery and precision have precluded the widespread adoption of these algorithms in comparison to the function-centric software, *antiSMASH*.

The most prominent challenge of USS algorithms is discriminating between gene clusters and loci that have shared synteny for other reasons. For example, regions near the centromere have shared synteny across relatively large phylogenetic distances because of a reduced rate of gene rearrangement compared to other genomic regions (57). Therefore, low USS thresholds can falsely designate regions with low rearrangement rates as clusters, while high thresholds can exclude recently evolved gene clusters with narrow distributions. Existing USS algorithms have attempted to address this problem in different ways. For example, *EvolClustDB* omits clusters smaller than five genes and imposes thresholds that remove recently evolved clusters using a global heuristic model of gene cluster size and shared synteny region similarity (74). *CO-OCCUR* detects shared synteny loci that contain genes with predetermined functions in order to predict gene clusters (28), which limits its scope to targeted searches.

Recently evolved, small clusters with low USS signal are detected by relaxing USS thresholds and increasing the accuracy of USS locus boundaries and phylogenetic distribution. Small clusters have relatively low USS signal because they need larger distributions to reach unexpectedness thresholds. Recently evolved clusters have small distributions, which also require low USS thresholds to identify. However, lowering USS thresholds may simultaneously increase false positives, so the distributions of shared synteny loci need to accurately recapitulate the distributions of the clusters they represent to minimize how much thresholds need to be lowered. *CLOCI* groups USS loci into homologous locus groups (HLGs) that recapitulate reference gene cluster distributions, including the relatively restricted distribution of the psilocybin gene cluster (43). We attribute the quality of HLG distributions to accurately defining shared synteny locus boundaries which allows for accurate locus-locus similarity calculations. *CLOCI* builds cluster boundaries from the most widely distributed combinations of gene families that underlay the cluster (Figure 3b). This approach is size-agnostic and detects small clusters such as the three gene penicillin cluster, with the caveat that two gene clusters are not part of initial unexpected synteny detection.

The signature of coordinated gene evolution can also help separate gene clusters from USS loci that are not clusters by increasing detection orthogonality. In *CLOCI, w*e implement four proxies of coordinated gene evolution to improve detection of true gene clusters from USS loci. These approximate selection for gene colocalization (TMD), the degree of gene coinheritance within an HLG/GCF (GCL), protein structural conservation (CSB), and loss and horizontal transfer (PDS).

Metabolic gene clusters that perform broadly distributed, or generalized, functions are identified within the highest TMD USS loci, whereas more specialized functions tend to have lower TMD values. The USS proxy is based on detecting unexpected TMD values of groups of co-occurring gene families, though a final TMD threshold is also applied to HLGs since multiple groups of shared synteny loci may represent a GCF. Increasing HLG TMD thresholds increasingly enriches general metabolic process genes. On the other hand, thresholds above 0.4 give diminishing returns for enriching secondary metabolic process genes, a subset of the general metabolic process category. This observation aligns with our finding that specialized metabolite biosynthesis clusters have the minimum observed TMDs, whereas the largest TMD clusters perform more essential metabolic functions (47) (Figure 2a). The limited distribution of specialized clusters may be attributed to their selection by impermanent, specific ecological factors, while clusters that perform essential/broadly distributed functions, such as biotin biosynthesis and iron assimilation, are less affected by environmental heterogeneity (87–90).

Gene Commitment to the Locus (GCL) quantifies the similarity between a gene’s distribution and the distribution of its HLG, as a proxy for the degree to which genes are co-inherited within gene clusters. Selection is thought to maintain the clustered state through selection against partial loss of co-adapted genes contributing to the same phenotype (91) and through increased fitness of clustered genes following horizontal transfer (51). We therefore expect clustered genes to be more committed to their clusters than non-clustered genes. GCL enriches secondary metabolic process genes across the tested thresholding range, which suggests genes within specialized gene clusters more often remain clustered. This may result from selection acting on the whole specialized function, which preserves the clustered state by increasing the fitness of clustered genes (73,92). GCL may have diminishing returns in filtering for general metabolic gene clusters because essential metabolic functions are likely to be retained by selection even in the absence of clustering in certain lineages. For instance, lineages may have insufficient effective population sizes to drive clustering through selection for metabolic efficiency (23,24,93).

We implemented the Conservative Substitution Bias (CSB) as a proxy for selection that maintains the coordinated function of genes in gene clusters (93). Genes within clusters coordinately carry-out cooperative metabolic function by co-locating, co-expressing, and binding (93–95). While coordinated function is necessary for clusters to produce their phenotypes, selection for coordinated function may additionally be influenced by the accumulation of toxic intermediate metabolites that are produced from partial metabolic pathways that result from discordance (52). Filtering using CSB thresholds may be effective in enriching specialized metabolic clusters in part due to CSB from selection against toxic specialized metabolic phenotypes and intermediate metabolites (52).

Phylogenetic Distribution Sparsity (PDS) is the most powerful filtration proxy for secondary metabolic gene clusters, even though many have a low PDS value. PDS approximates the prevalence of horizontal transfer and loss that shape the distribution of metabolic gene clusters (24,51,55,90). Gene clusters may be subject to elevated horizontal transfer when they contain complete selectable phenotypes (51,90,92) and can be rapidly lost in lineages when the cost of the cluster phenotype exceeds its fitness contribution (55). Additionally, the prevalence of horizontal transfer and loss may vary by lineage (92), or type of cluster (Figure 2a), which results in the hypervariability of PDS values (minimum 0%, median 81.3%). Gene clusters with specialized functions are often tied to environment-interaction phenotypes, and therefore selected by the ecology of the organisms that contain them (30,31,43,73). This specialization may lead to horizontal transfer among organisms occupying similar niches (43) and cluster loss upon niche switching. Our results support PDS as a proxy for specialized metabolic clusters, as increasing minimum PDS thresholds superlinearly extracts clusters with secondary metabolic process genes. PDS also enriches general metabolic process genes, perhaps in part due to spurious annotation of cryptic specialized metabolic functions as general metabolic processes (96), though PDS selectivity for specialized metabolite clusters becomes more prominent as PDS surpasses 0.2. Horizontal transfer can act across large phylogenetic distances relative to the phylogenetic distribution of the cluster and thus drastically increase specialized cluster PDS signal compared to loss (31,72). Some accessory metabolic clusters that perform functions with broad usefulness, such as nitrate assimilation, may also transfer across large phylogenetic distances (25,97). However, PDS for the nitrate assimilation cluster is not as high as specialized metabolite clusters with smaller transfer distances (72) because nitrate assimilation is such a broadly distributed accessory function (Figure 2a). It is important to note that in situations where TMD approaches a complete distribution, any decrease in PDS could potentially be attributed to sampling limitations (Equation 4). However, we did not identify any reference clusters or extracted GCFs that had nearly complete distributions.

Proxies for coordinated gene evolution are partially linearly correlated. We found that TMD is correlated with the other proxies (adjusted R^2^ > 0.5), which suggests that the distribution of a cluster positively associates with the variance in selection for coinheritance, coordinated function, and prevalence of horizontal transfer and loss. The correlation between TMD and CSB is relatively high (adjusted R^2^ = 0.833), indicating that broadly distributed clusters have increased selection for structural conservation. This may be because clusters that have large TMDs often perform essential metabolic functions that are under intense purifying selection to preserve the integrated function, whereas clusters that perform more specialized functions can adapt to new ecological roles and experience positive selection (80–82). Alternatively, the correlation between TMD and CSB may be result from increased signal from conservative amino acid substitutions at greater phylogenetic distances. Interestingly, enrichment using minimum thresholds for CSB mirror the results for GCL (Figure 2), though the congruence between GCL and CSB is not completely explained by collinearity between selection for coinheritance and coordination because these proxies are not completely correlated (adjusted R^2^ = 0.446).

### *CLOCI* more accurately identifies gene cluster families when compared to existing algorithms

When compared to the function-centric algorithm *antiSMASH, CLOCI* recovers more biosynthetic gene clusters and greater cluster diversity. The 7.35% of clusters *CLOCI* fails to recover can largely be attributed to sparse sampling in our dataset. *CLOCI* also has greater general cluster recovery when compared to an existing USS algorithm, *EvolClust.* We attribute *CLOCI* improvements to USS detection by more precisely inferring cluster boundaries through a robust gene cluster grouping algorithm that globally identifies GCFs. The improvements to recovery and boundary accuracy position *CLOCI* as a paradigm-shift in gene cluster detection from function-centric algorithms to coordinate gene evolution detection.

*CLOCI* recovers more gene clusters than function-centric algorithms. *CLOCI* and the function-centric algorithm, *antiSMASH,* both detected the majority of biosynthetic gene clusters in the dataset (91.2% and 76.1% respectively). *CLOCI* additionally inferred 100% of the non-biosynthetic gene cluster categories because USS among gene clusters is a property independent of gene function. In contrast, *antiSMASH* was initially designed to detect biosynthetic gene clusters, and the capacity for *antiSMASH* to infer diverse metabolic pathways can be attributed to the versatility of the modeled core genes it searches for. Modeling canonical biosynthetic clusters, such as polyketide and nonribosomal peptide clusters, can reveal most of the known fungal biosynthetic cluster diversity because these clusters can synthesize diverse metabolites from modular core genes (87,88). Non-biosynthetic clusters can be incorporated into *antiSMASH* detection, though function-centric non-biosynthetic screening may be relatively restricted to the modeled cluster family (27) because non-biosynthetic cluster classes have relatively conserved phenotypes (55). Similar to non-biosynthetic functions, noncanonical biosynthetic gene clusters, such as the psilocybin cluster, may have discrete distributions with conserved phenotypes, so modeling these functions may also have limited capacity for inferring new clusters. In other words, noncanonical clusters may not contain versatile, modular core genes and may have limited capacity for diversified biosynthesis. Therefore, even if function-centric detection continues to add newly discovered core genes it cannot infer cluster classes *de nov*o because they have not been modeled. *CLOCI* inherently detects these GCFs without having to account for their functions *a priori*.

While *CLOCI* successfully identifies most reference clusters, five of the 68 reference gene clusters (7.35%) were not detected primarily due to genome sample limitations. We failed to recover the zearalenone cluster because it is minimally sampled and has a small distribution within *Fusarium* (98), which did not meet USS thresholds. The mycophenolic acid gene cluster was overlooked because the locus was incorrectly assembled due to low support that is partially attributed to excluding *Penicillium brevicompactum* genomes that failed our genome quality control prior to the analysis. Interestingly, neither *antiSMASH* nor *CLOCI* both detected the cephalosporin gene cluster in *Acremonium chrysogenum,* perhaps because it has a bacterial origin (99) and thus is not supported from our fungal dataset. Both algorithms also failed to detect the ent-kaurene cluster, which *CLOCI* misses because it primarily comprises multiple tandem gene duplications. Recent tandem duplicates are binned into the same homology group and our *CLOCI* analysis settings constrained detection to at least three unique homology groups within a five gene sliding window. Missing clusters highlight the benefit of further genome sampling, and suggest *CLOCI* recovery will improve with further tuning of algorithm variables.

*CLOCI* improves USS-based detection by implementing an approach that more accurately defines shared synteny loci (Figure 3). The discrepancy in recovery of our cluster recovery dataset between *CLOCI* (92.7%) and the existing USS algorithm, *EvolClust* (51.0%), may be partially explained by our increased genome sample because both algorithms rely on sufficiently sampling loci to support cluster detection. However, the *EvolClustDB* also recovers fewer clusters (75.0%) than *CLOCI* (91.0%) in the independent cluster boundary dataset (Supplementary Table 5). The cluster boundary dataset is primarily composed of *Aspergillus* spp., a genus which is well-sampled by both algorithms, and thus we attribute the discrepancy in recovery to the alternative approaches in inferring shared synteny loci. *EvolClust* implements a global heuristic model of shared synteny locus sizes with a minimum size of five genes to mitigate false positive gene cluster designation. In contrast, *CLOCI* can detect clusters from as small as two genes up to the chromosome length by assembling gene clusters from domains of shared microsynteny that more accurately recapitulate the boundaries of homologous loci. *CLOCI* separates the three gene *Ustilago* nitrate assimilation and *Saccharomyces* galactose metabolism gene clusters from the surrounding unexpectedly conserved regions, whereas *EvolClust* infers both gene clusters with 16 extra genes each.

Our results corroborate previous findings (54) that shared synteny detection more precisely infers cluster boundaries than both standalone boundary prediction in *antiSMASH* and *antiSMASH* with *CASSIS* (Figure 3). According to our functional cluster boundary definition, boundary inference using *antiSMASH* with or without *CASSIS* on average infers more than double the reference cluster size. *CASSIS* overestimating cluster boundaries can be attributed to its algorithm operationally defining gene clusters boundaries as entire coregulated regions that may include multiple clusters. Indeed the coregulated fumagillin and pseurotin clusters in the reference dataset are fused in *antiSMASH* output with and without *CASSIS* (47,100). In contrast, *CLOCI* predicts a median zero extraneous and missing gene count per reference cluster and separately reports the fumagillin and pseurotin clusters. While the majority of *CLOCI* clusters are reported as a single shared microsynteny domain (65.6% of reference clusters), we identified some reference clusters as two domains (27.6%) and one cluster as three domains. These domains are directly adjacent, though *CLOCI* does not report them with other evidence of linkage. The granularity of sub-cluster domains is tunable, and poorly sampled clusters will become reportable as a single domain as the quality of the genome sample improves.

### *CLOCI* enables unbiased gene cluster analysis

*CLOCI* does not depend on *a priori* assumptions of function, which enables unbiased comparative ecological studies on genomes’ cluster repertoires that include ecologically-relevant non-biosynthetic clusters. *CLOCI* further extends the capacity to globally detect gene cluster categories to GCF circumscription, where GCFs are globally identified using Markov Clustering (MCL) and the granularity of GCF identification is tunable through a single parameter. Examining the network topology of these GCFs can provide insight into the evolution of gene clusters, including evidence of horizontal transfer. We also sift these GCFs for unreported primary and accessory metabolic gene clusters by filtering minimum criteria for coordinate gene evolution proxy values.

*CLOCI* presents an unbiased framework for ascertaining the ecological functions of genomes as products of their gene cluster repertoires. Gene cluster detection algorithms are extensively implemented to study and compare the ecology of genomes. However, these studies primarily implement function-centric algorithms that are biased toward well-studied lineages and only detect canonical specialized metabolite biosynthesis gene clusters (39,40). Despite this bias, function-centric detection has dominated the literature, in part because previous function-agnostic algorithms do not recover as many gene clusters (Figure 3a). *CLOCI* recovers more gene clusters without *a priori* assumptions of function, which presents an unbiased framework for comparatively profiling gene cluster repertoires. This approach has the advantage of cataloging ecologically-significant non-biosynthetic gene clusters that are overlooked by function-centric algorithms, such as nitrate assimilation and galactose metabolism, which presents a broader window into the ecologies of gene cluster repertoires.

*CLOCI* implements a globally tunable framework for circumscribing GCFs. Robust GCF circumscription accounts for microsynteny, amino acid similarity, and protein domain presence-absence to group gene clusters into ecologically-significant GCFs (30,31), and tie biosynthetic GCF granularity to metabolite structural modifications (101–103). GCF circumscription algorithms have had to be tuned for each type of gene cluster because these algorithms use weighted coefficients that must be tailored to the different evolutionary rates of different GCF types (28,104). In contrast, *CLOCI* implements a globally-tunable model that affects granularity for all types of gene clusters using a single parameter, the MCL inflation value. The MCL random-walk approach to graph clustering is attractive for GCF identification because MCL incorporates normalization at each step, which allows for divergent local graph similarities that reflects the different evolutionary rates of different GCFs. The size-flexibility of this approach is demonstrated by *CLOCI* inferring the nitrate assimilation GCFs (assimilative) in multiple phyla, whereas the psilocybin GCF (noncanonical biosynthetic) was recovered in only five organisms. Increasing the inflation value will globally increase the granularity of GCF inferences, which can be compared with metabolic data to identify functionally-significant modifications in GCFs.

GCF network topology is consistent with horizontal transfer events and deep phylogenetic divergence in the evolution of gene clusters. We identified at least five nitrate assimilation GCFs that contain characterized nitrate assimilation genes (18,25,105). The locus-locus network topology suggests these divergent GCFs are homologous (Figure 4f). The GCFs correspond with discrete subnetworks because deep phylogenetic divergence results in low sequence similarity and variation in gene composition (25,82,97). For example, some Saccharomycotina nitrate assimilation clusters contain unique transcription factors that result in decreased locus-locus similarity with other nitrate assimilation GCFs (81). Alternatively, convergent evolution may also result in multiple GCFs. Horizontal transfer is suggested by the presence of distantly related taxa within a subnetwork. For example, the aggregation of Hypocreaceae*/*Bionectriaceae (Ascomycota) and Basidiomycota nitrate assimilation clusters in a common GCF is consistent with known horizontal transfer between these lineages (25). We also identified a nitrate assimilation GCF composed of Mucoromycota and Basidiomycota clusters. Phylogenetic analysis of all genes within this GCF supports the cluster was transferred from Mucoromycota species to *Amanita* (Basidiomycota). The cluster was only recovered in ectomycorrhizal *Amanita* spp. (83) and is missing from the saprobic *Amanita inopinata* and *Amanita thiersii. T*his suggests acquisition of the nitrate assimilation GCF coincided with the emergence of the mycorrhizal ecology in *Amanita* species, possibly selected by the increased nitrogen demand imposed by host trees.

Proxies of coordinated gene evolution characterize the evolutionary properties of GCFs and can select for clusters of interest. The vast majority of *CLOCI* output are not yet characterized gene clusters (Figure 3c). The quantity of uncharacterized clusters within our results is consistent with the hypothesis that most clusters have not been studied (4,106,107), and presents an exciting opportunity for mining novel biosynthesis and ecologically important clusters from *CLOCI* output. To demonstrate *CLOCI’s* capacity to identify generalized cluster functions, we identified an unreported gene cluster with high log normalized TMD that contains the basidioferrin synthetase throughout the mushroom-forming subphylum, Agaricomycotina. If the function of this cluster is conserved, then it presents a possible conserved mechanism for iron assimilation in Agaricomycotina. We additionally searched the upper 99 percentile of PDS scores to identify putative specialized metabolite clusters. We identified a GCF with an undescribed nonribosomal peptide biosynthetic cluster (Figure 4b) in three different taxonomic classes, which could potentially be explained by multiple horizontal transfer events or convergent origins. The approach we employed to identify this putative specialized metabolite gene cluster can be used as the foundation of specialized metabolite biosynthesis detection, including relating hypothesized noncanonical biosynthetic pathways to *CLOCI-*predicted clusters.

### *CLOCI* is a lineage-independent foundation for detecting shared synteny loci that are not clusters

*CLOCI* is tailored toward gene cluster detection in fungi, though the algorithm can be applied to other kingdoms. It is expected that the same principles will be adaptable for detecting accessory gene clusters in prokaryotes because prokaryotic clusters often comprise multiple discrete gene families. However, clusters that exclusively comprise recent homologs, particularly due to tandem gene duplication, will not be detected by *CLOCI*. Tandem duplicate clusters are common in animals and plants. Future work will focus on expanding the capacity to detect such clusters by increasing the sliding window, accounting for tandem duplications in homology group co-occurrences, separating tandem duplicate orthologs into independent homology groups, and further tuning detection parameters with machine learning.

*CLOCI* can be implemented to classify HLGs and identify unexpectedly shared synteny regions without proxies, or proxies can be adapted to different types of genomic loci that are also under selection for coordinated gene evolution. On its own, HLG inference is an algorithmic adaptation of orthologous gene group (orthogroup) circumscription to the locus level, which thus provides a means for comparing the presence and absence of functionally significant gene neighborhoods. *CLOCI* HLG detection can even predict loci that are fragmented across multiple contigs, which can be used to inform genome assembly scaffolding (108). Following HLG inference, proxies of coordinated gene evolution can be tuned to accommodate different types of shared synteny loci. For example, bacterial defense islands contain genes and gene elements necessary for recognition and immunity against viruses and other antagonistic elements (109) and retroviral DNA is integrated into host genomes in discrete regions (110). *CLOCI* HLGs provide a powerful foundation for identifying these shared synteny loci in conjunction with gene function assignment.

### Recommendations and considerations for implementation

*CLOCI* is constructed in a comparative genomic framework, and thus requires a large enough sample of sufficiently diverse genomes to identify unexpectedly shared synteny. The inputted genome sample should be based on lineages’ rates of microsynteny decay and the distribution of their gene clusters. *CLOCI* may best be implemented at least on a subphylum-level dataset to account for the majority of horizontal transfers.

Improvements to *CLOCI’s* memory usage and time complexity should be focused on homology group combination inference, CSB quantification, and GCF circumscription, which involve pairwise loci comparisons and self-alignment of entire homology groups. We implement methods to decrease the amount of comparisons, such as homology group pair seeding and gene cluster clan classification; however, the time complexity of these modules remains non-linear. A compiled language implementation of these modules would additionally increase their throughput.

## DATA AVAILABILITY

*The data underlying this article are available in the article and in its online supplementary material. CLOCI is available at gitlab.com/xonq/cloci. Mycotools is available at gitlab.com/xonq/mycotools*.

## SUPPLEMENTARY DATA

Supplementary Data are available at bioRxiv.

## ACKNOWLEDGEMENTS

We would like to acknowledge the Ohio Supercomputer Center for providing cutting-edge high performance computing resources. We thank Dr. Emile Gluck-Thaler for consultation regarding the development of *CO-OCCUR* and its application as the conceptual foundation of *CLOCI*.

## GLOSSARY

CSB: Conservative Substitution Bias, a proxy for selection for structurally conservative amino acid substitutions
CLOCI: Co-occurrence Locus and Orthologous Cluster Identifier, a novel function-agnostic gene cluster detection algorithm
GCF: gene cluster family or group of homologous clusters, a subset of HLGs
GCL: Gene Commitment to the Locus, a measurement of the degree to which a gene family is committed a particular locus
HG: homology group, or group of homologous genes
HLG: homologous locus group consisting of loci with recent shared decent and unexpectedly shared synteny
MCL: Markov clustering, a graph clustering algorithm that implements a random-walk approach to aggregating similar graph vertices
MMI: Minimum Mean Identity, a measurement of the average minimum alignment percent identity between homologous genes within HLGs
MMP: Minimum Mean Positives, a measurement of the average minimum alignment percent positive substitutions between homologous genes within HLGs
PDS: Phylogenetic Distribution Sparsity, a measurement of the sparsity of a trait’s distribution
TMD: Total Microsynteny Distance, a measurement of the microsynteny tree branch length a trait is distributed

