## supplementary figures for "*CLOCI:* Unveiling cryptic gene clusters with generalized detection"

Supplementary Figure 1: Visualization and linear correlation analysis via ordinary least squares regression between HLG coordinate gene evolution proxies; TMD is log normalized. TMD v GCL Adj. R-squared 0.606, p < 0.001; TMD v PDS Adj. R-squared 0.542 p < 0.001; TMD v CSB Adj. R-squared 0.833, p < 0.001; GCL v PDS Adj. R-squared 0.127, p < 0.001; GCL v CSB Adj. R-squared 0.446, p < 0.001; PDS v CSB Adj. R-squared 0.509, p < 0.001

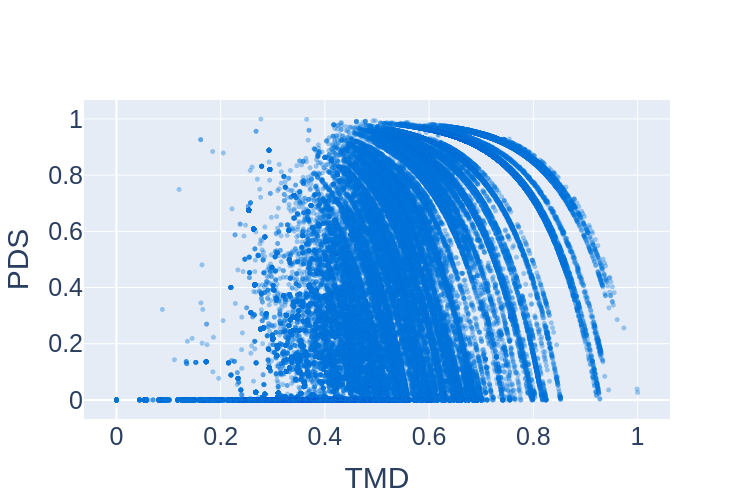

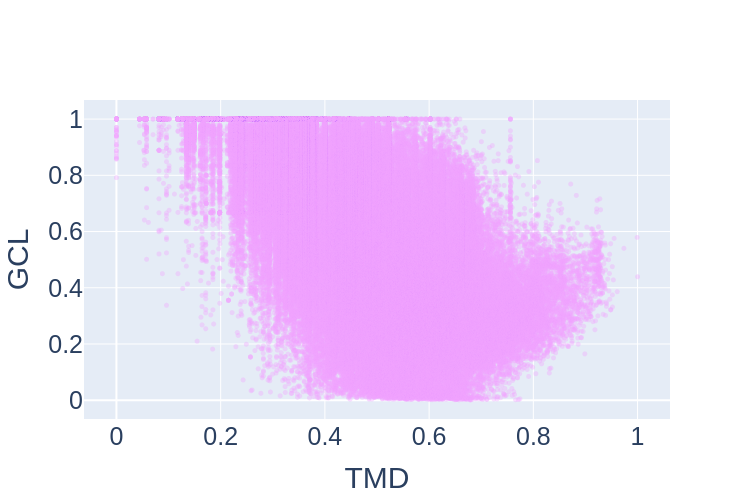

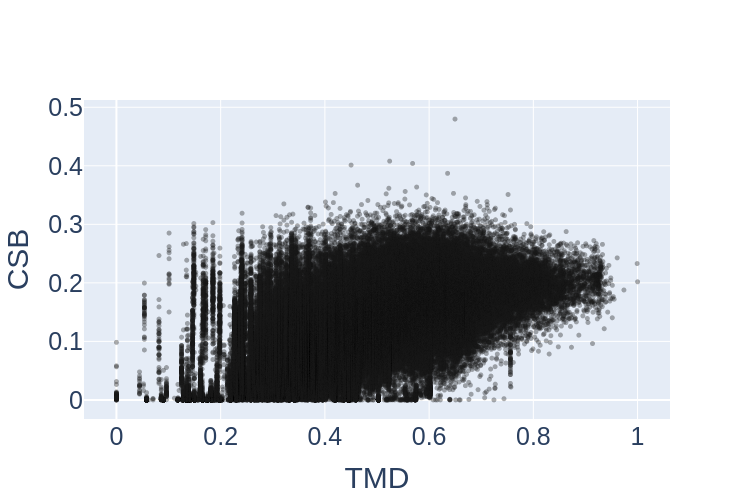

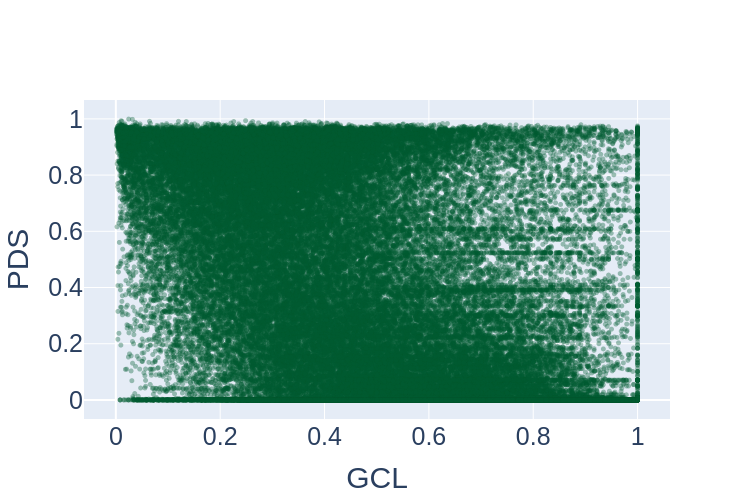

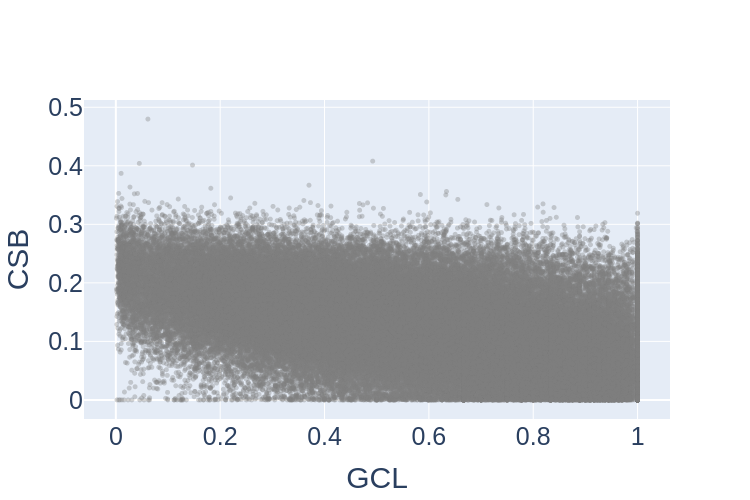

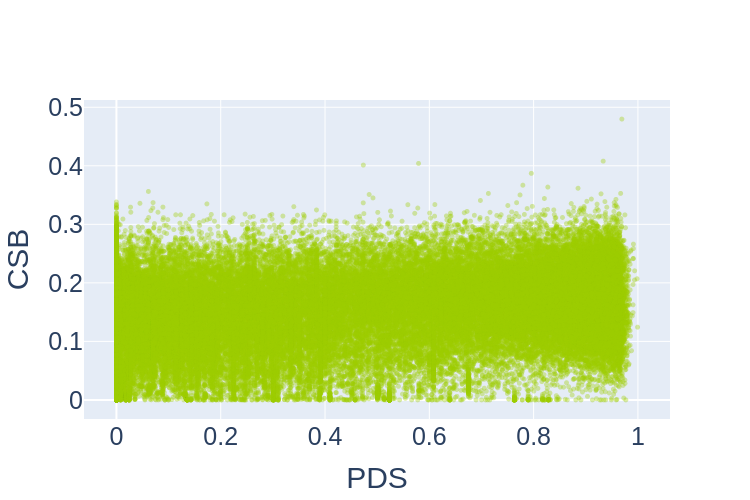

Supplementary Figure 2: Sunburst charts representing distribution of MycotoolsDB taxa (top) inputted into *CLOCI* analysis and distribution of MiBIG taxa (bottom)

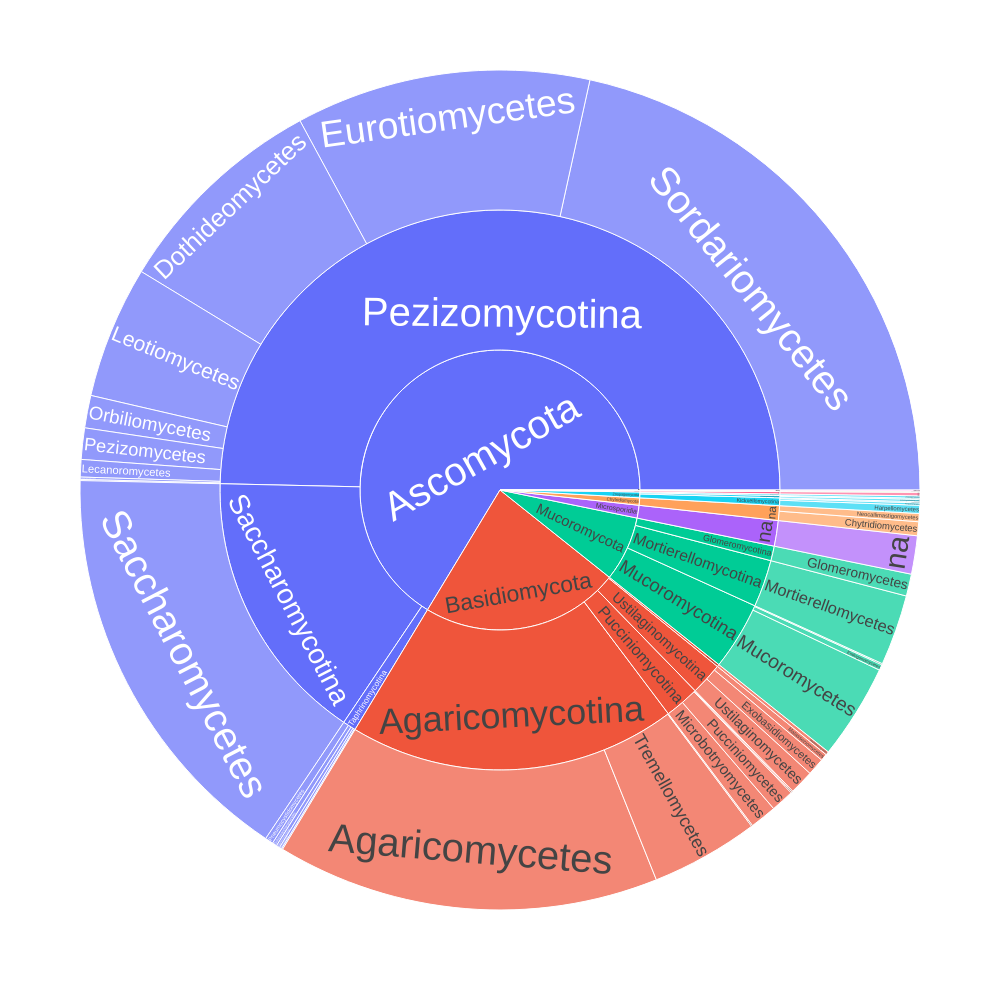

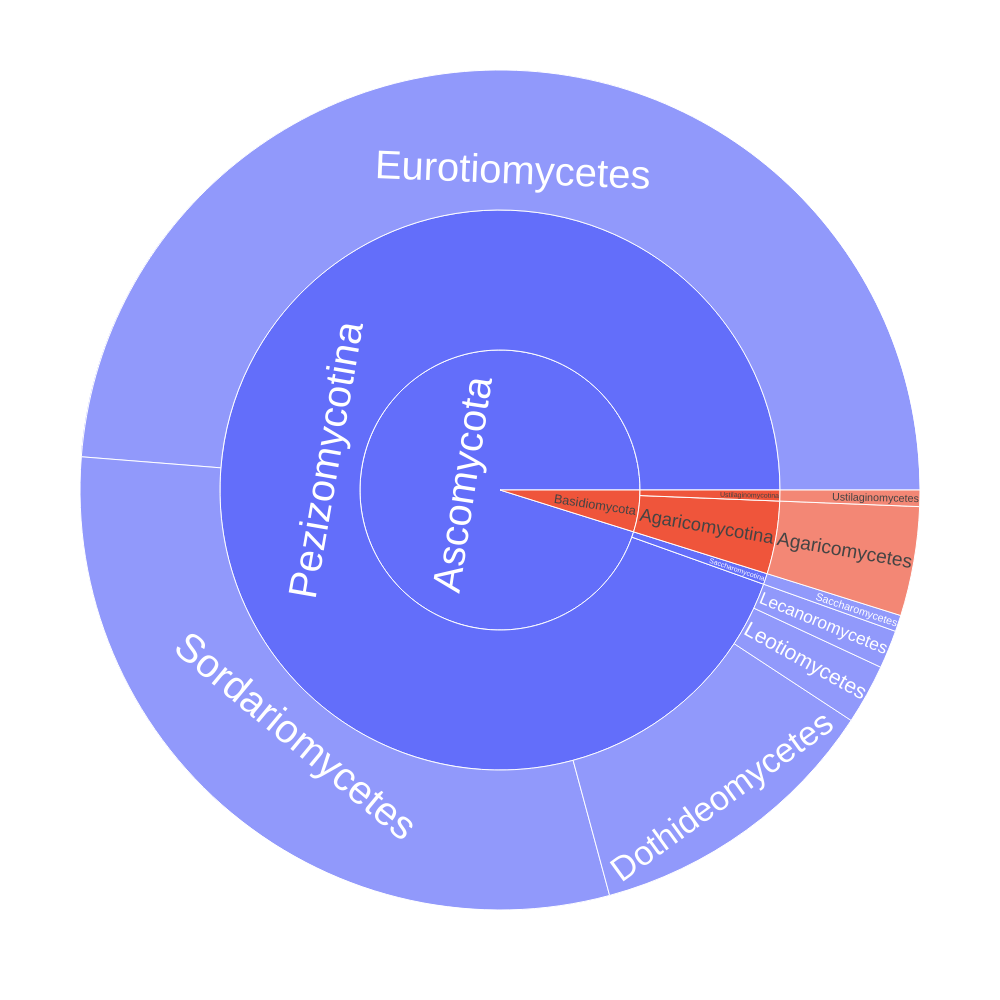
